# Identification of alkylsalicylic acids in Lentisk oil (*Pistacia lentiscus* L.) and cytotoxicity on Human Normal Dermal Fibroblasts

**DOI:** 10.1101/2020.11.17.373142

**Authors:** Nabiha Benalia, Abdenour Boumechhour, Sergio Ortiz, Cristian A. Echague, Thorsten Rose, Bernd L. Fiebich, Smain Chemat, Sylvie Michel, Brigitte Deguin, Saliha Dahamna, Sabrina Boutefnouchet

## Abstract

*Pistacia lentiscus* L. (Anacardiaceae) is widely distributed in the Mediterranean basin. Its fruit oil is used in traditional medicine to treat burns, skin impairments as well as inflammatory diseases as soothing massage or internal use. An increased interest is spotted lately with several commercial brands are spun portraying the benefits of this oil but with no stringent regulations to ascertain its safe use as an edible or cosmeceutical product. This work concerned the investigation of secondary metabolites presents in *Pistacia lentiscus* fruits oil using both GC-MS and HPLC-DAD-MS technics, and the evaluation of cytotoxicity on human normal dermal fibroblasts to assess safety of use as cosmetic ingredient. This study stands as the first one to report the identification of alkylsalicylic acids in fruits oil and unsaponifiable fraction of *Pistacia lentiscus* fruit oil which calls for therefore, quantification of alkylsalicylic acids, known as skin irritants, in *Pistacia lentiscus* oil, used as nutraceuticals or cosmeceuticals by manufacturers.

## INTRODUCTION

Lentisk, *Pistacia lentiscus* L is an evergreen shrub or tree from Anacardiaceae family, largely known as Darou, dherou or Drou in North Africa, Listincu or Chessa in Sardinia; or Mastiha tree in Greece. Its repartition covers all the Mediterranean area, from the Iberic peninsula to the Middle East. Mastic is the oleoresin obtained from the tree after incision of its trunk and is produced in abundance by the *Chia* variety occurring in the Greek Chios island. Despite the large use of mastic for medicinal or dietary purpose (Pachi et al, 2020), the application of vegetal oil obtained from fruits is still less known.

An archaeological study from the eastern Mediterranean brought evidence of fruit oil production via fruits’ squeezing in Roman and high medieval sites in Sardinia island and Corsica. The same processing method exists also traditionally in eastern Algeria and Tunisia, but not described in the western part of Mediterranean, which may indicate the probable influence of Roman empire colonization. Lentisk oil was mainly used for lighting, treating burns and wounds, and as food dressing (Lanfranchi et al, 1999; Lanfranchi and Bui, 1998). Several reports indicated the limited geographical distribution of lentisk oil users with predominance use is linked to traditional pharmacopoeia of eastern and central parts of Algeria and Tunisia for the treatment of skin, respiratory conditions and rheumatism. According to Djerrou et al. study, the use of *Pistacia lentiscus* fixed oil reduced the inflammatory phase, stimulated wound contraction and reduced the epithelization period to those treated with pharmaceutical grade oitment Madecassol® (Dierrou et al, 2007). Mammeri at al. combined it with honey and confirmed its superiority to accelerate wound healing via contraction compared to a commercial skin protector cicatryl© (Maameri et al, 2012).

Chemical composition of *Pistacia lentiscus* oil has been mainly focusing on saponifiable fraction of the oil including fatty acids, phytosterols and tocopherols (Charef et al, 2008; Mezni et al, 2012; Trabelsi et al, 2012). However, an increased interest is spotted lately with several commercial brands are spun portraying the benefits of this oil but with no stringent regulations are in force to ascertain its safe use as an edible or cosmeceutical product. This emerging popularity calls for the establishment of quality indicators to ensure quality of marketed oils, reduce the risks of adulteration and/or misuse and ascertain its safe use.

Thus, in this study, we focused in Lentisk oil with an objective of two folds, the first one consists to compare fourteen oil samples in order to depict quality indicators by spotting differences in secondary metabolites profiles using both GC-MS and HPLC-DAD-MS technics, the second one was to assess cytotoxicity on human normal fibroblast to elaborate its potential safe use in dermatological applications.

## 1. MATERIALS AND METHODS

### 1.1. Oil samples and reagents

Artisanal samples were obtained from rural families located in the area of Jijel (Sidi abdelaziz, Settara, Ouled rabeh), Blida and Elkala (Eltaref) (Supporting informations). Artisanal production consists in crushing of mature fruits collected during the period of end of autumn-beginning of winter, then submitted for maceration in cold water. In certain cases, the mixture may be heated to increase oil’s yield. After filtration, the oil is separated from water by decantation to afford a yellow green viscous liquid. The obtained yields ranged between 16% and 19% (Lanfranchi and Bui, 1998). In parallel, semi-artisanal samples were obtained from two cooperatives: Ladjoudane (Akbou) and Bouannani (Jijel). These oils are prepared following the same process described earlier but extraction is realized using a hydraulic press equipped with fibre disks called “scourtins”. In another part, six commercial samples were purchased in the same areas (Table 1 and 2). Methanol (MeOH) HPLC grade, dimethylsulfoxide (DMSO), ethanol (EtOH), gallic acid, quercetin, Folin-Ciocalteu reagent, AlCl_3_·6H_2_O, Na_2_CO_3_, CH_3_COONa, BF_3_ (4%)-methanol, hexane, were purchased from Sigma Aldrich (St. Louis, USA).

**Table 1.** Fatty acids composition of artisanal *Pistacia lentiscus* L. oil

|  | 1 | 2 | 3 | 4 | 5 | 6 | 7 | 8 |  |
| --- | --- | --- | --- | --- | --- | --- | --- | --- | --- |
| Artisanal samples | Bouanani | Akbou<br>2015 | Akbou<br>2017 | Settara | Ouled<br>Rabah | Sidi<br>Abdelaziz | El Kala | Blida | Mean+/-SD |
| % Fatty acids |  |  |  |  |  |  |  |  |  |
| Palmitoleic acid 16 :1 | 2.38 | 1.72 | 2.07 | 3.03 | 2.02 | 2.42 | 2.1 | 2.76 | 2.31+/-0.43 |
| Palmitic acid 16 :0 | 31.45 | 23.94 | 30.34 | 34.22 | 27.15 | 28.09 | 30.54 | 37.08 | 30.35+/-4.10 |
| *n*- Hexadecanoic acid 16:0 | 0.19 | 0.36 | 0 | 0 | 0.17 | 0.17 | 0 | 0 | 0.11+/-0.13 |
| Linoleic acid 18 :2 | 15.98 | 21.33 | 17.15 | 15.98 | 14.52 | 15.24 | 16.87 | 15.98 | 16.63+/-2.07 |
| Oleic acid 18 :1 | 39.67 | 46.33 | 42.83 | 43.96 | 52.84 | 51.89 | 48.6 | 41.12 | 45.91+/-4.88 |
| Stearic acid 18 :0 | 6.12 | 2.73 | 6.63 | 2.19 | 2.2 | 1.11 | 1.15 | 1.87 | 3.00+/-2.16 |
| Saturated Fatty acids SFA | 37.76 | 27.03 | 36.97 | 36.41 | 29.52 | 29.37 | 31.69 | 38.95 | - |
| Unsaturated Fatty acids<br>UFA | 58.03 | 69.38 | 62.05 | 62.97 | 69.38 | 69.55 | 67.57 | 59.86 | - |
| UFA/SFA | 1.54 | 2.57 | 1.68 | 1.73 | 2.35 | 2.37 | 2.13 | 1.54 | - |

**Table 2.** Fatty acids composition of commercial *Pistacia lentiscus* L. oil

|  | 9 | 10 | 11 | 12 | 13 | 14 |  |
| --- | --- | --- | --- | --- | --- | --- | --- |
| Commercial samples | El Wafia | Zazia | Belkis | Zahrat el Atibaa | Al Fourssan | Sultane | Mean+/-SD |
| % Fatty acids |  |  |  |  |  |  |  |
| Palmitoleic acid 16 :1 | 0.13 | 0 | 2.03 | 1.69 | 2.31 | 1.28 | 1.24+/-0.97 |
| Palmitic acid 16 :0 | 16.41 | 16.25 | 24.78 | 31.58 | 28.42 | 22.35 | 23.3+/-6.25 |
| *n*- Hexadecanoic acid 16:0 | 0 | 0 | 0 | 0 | 0 | 0 | 0 |
| Linoleic acid 18 :2 | 47.46 | 44.95 | 15.46 | 17.37 | 15.35 | 20.69 | 26.88+/-15.11 |
| Oleic acid 18 :1 | 28.72 | 31.35 | 54.62 | 47.58 | 50.13 | 53.5 | 44.32+/-11.37 |
| Stearic acid 18 :0 | 5.28 | 5.16 | 2.14 | 1.3 | 2.15 | 1.84 | 2.98+/-1.76 |
| Saturated Fatty acids SFA | 21.69 | 21.41 | 26.92 | 32.88 | 30.57 | 24.19 | - |
| Unsaturated Fatty acids UFA | 76.31 | 76.3 | 72.11 | 66.64 | 67.79 | 75.47 | - |
| UFA/SFA | 3.52 | 3.56 | 2.68 | 2.03 | 2.22 | 3.12 | - |

### 1.2. Pistacia lentiscus L. oil composition

Fatty acids are analyzed after transesterification using BF_3_ into their corresponding methyl-esters (Charef et al, 2008; Mezni et al, 2012; Trabelsi et al, 2012). Briefly, 50 mg of oil are diluted in 1 mL hexane, to which 0.5 mL of BF3 (14%)-methanol is added. The mixture is kept at 60°C during 30 minutes after which 1 mL of water added. Then, the organic phase containing the FAME mixture is separated for GC-MS and GC-FID analysis. Analyses are performed using GCMS-QP2010 Ultra (Shimadzu Co. Kyoto. Japan) equipped with a fused silica capillary column (Rtx-5MS; 30 m × 0.25 mm inner diameter, film thickness 0.25 μm, Thames Restek. UK), and an Agilent 5973N MS detector. GC-FID was performed with a Master GC-Dani, France, equipped with a HP5 column (5% Phenyl-Hexyl, 100 m, internal diameter 0.25 mm, thickness 0.2 μm) and a FID detector.

The following analytical conditions are used : oven temperature is programmed to start at 140°C, hold for 1 min, then increased to 200°C at a rate of 5°C/min, then hold for 3 min, then increased to 215°C (rate 5°C/min, hold for 5 min), then increased to 240°C at a rate of 10°C/min, and finally hold for 10.5 min. Injector temperature is set at 270 °C; carrier gas: helium, flow, 0.95 mL/min; splitting ratio 1:20; injection volume: 1 μL; interface temperature: 240 °C; while for the mass spectrometry interface, the MS source temperature is set at 220 °C with an ionization energy of 70 eV. For other volatiles analysis, the oven temperature is programmed at 70°C, hold for 5 min, then increased to 120°C (rate 5°C/min, hold for 2 min), then increased to 180°C (rate 30°C/min, hold on 12 min), and finally increased to 270°C (rate 30°C/min, and kept for 2 min. Fatty acids were identified by comparison of their recorded mass spectra with the NIST14 library and the calculated retention indices (RI) of corresponding FAME fatty acid methyl esters.

### 1.3. Analysis of unsaponifiable fraction

The recovery of unsaponifiable fraction for artisanal samples 2, 7, and 8 is conducted following the procedure described in AFNOR NF T 60-206. Briefly, 50 mL of 2N ethanolic solution of KOH are added to 5 g of oil. The mixture is then refluxed for one hour. After evaporation, 50 mL of water is added, the suspension is extracted three times with 100mL diethyl ether (3×100 mL), washed with aqueous KOH (0.5 N) followed by water, then dried on anhydrous Na2SO4 and evaporated under vacuum. Then, derivatization into silyl esters is performed. Briefly, 5 mg of unsaponifiable fraction are placed with 0.5 mL of pyridine in a 2 mL vial. Then, 0.1 mL of hexamethyldisilazane (HMDS) and 0.04 mL of trimethylchlorosilane (TMCS) are added and the reaction mixture is mixed using a vortex then centrifuged. From the supernatant of the silylated mixture, 1 µL is directly submitted to GC-MS analysis. In this case, carrier gas is H2 with a flowrate of 1 mL/min and a split 1:20, oven is programmed increasing from 180°C to 270°C at 8°C/min with a hold at initial and final temperatures of 1 and 65 min respectively (Jasmica, 2001). MS Interface temperature is set at 240 °C while MS source temperature is set at 220 °C with an ionization energy of 70 eV. The injection volume was 1 µL.

### 1.4. HPLC-DAD-MS identification of alkylsalicylic acids in Pistacia lentiscus oil

Oil samples are analyzed on a HPLC-DAD-MS Thermo Scientific Dionex U3000 (Thermo-Dionex, Les Ulis, France) consisted of a quaternary pump (LPG-3400 SD), a thermostated autosampler (WPS-3000TSL), a thermostated column (TCC-3000SD), and a diode array detector (DAD-3000) on line with a quadrupole mass spectrometer (Surveyor MSQ plus System (Thermo-Dionex, Les Ulis, France). All oil samples were diluted in methanol (10 mg/mL). Solutions are filtered before injection on UptiDisc 0.45 M nylon filters (Interchim, Montluçon, France). 20 µL of each solution are injected and chromatograms are recorded at 210, 280, and 320 nm. Oven temperature is set at 30°C and the analysis is performed using a gradient elution: A (H2O, 0,5% formic acid) and B (ACN, 0.5% formic acid) as follows: 5% of B (0–10 min, isocratic), 5% to 100% of B (10-40 min, linear gradient), 100% of B (40-60 min, isocratic), 100% to 5% of B (60-70 min, linear gradient), 5% of B (70-75 isocratic) the flow rate is fixed at 0.5 mL/min.

### 1.5. Alamar blue cell viability assay on Normal Human Dermal Fibroblasts

Normal Human Dermal Fibroblasts (NHDF) are cultured in Dulbecco’s modified Eagle medium (Sigma, UK) supplemented with 10% fetal calf serum (Biowest Ltd., UK), 2% l-glutamine (Sigma), 100 U/ml penicillin, (Sigma), and 100 g/ml streptomycin (Sigma). After counting the number of cells in a particle counter (Euro Diagnostics, Krefeld, Germany), NHDF are seeded in 96-well plates at a density of 8,000 cells/well and incubated at 37°C with 5% CO2 overnight. Cells are treated by 0.1, 1, 5, 10, 25, 50 and 100 µg/ml, of the extracts in 100 μL of media. Cells are incubated with sodium fluoride (NaF, 250 μg/mL), to induce cell death as the positive control, or with DMSO only (negative control). The experiment is conducted in quadruplicates. After 24 hours of incubation, 10 μL AlamarBlue® (Thermo Fisher Scientific, Waltham, MA, USA) is added to each well. After 2 hours, the fluorescence is measured with a fluorescence spectrophotometer (Polarstar, BMG, Offenbug) using 544EX nm/590EM nm filter settings. The amount of fluorescence is proportional to the number of living cells and corresponds to the cells’ metabolic activity. Damaged and nonviable cells have lower innate metabolic activity and thus generate a proportionally lower signal than healthy cells. The active ingredient of AlamarBlue® (resazurin) is a nontoxic, cell permeable compound that is blue in color and virtually nonfluorescent. Upon entering cells, resazurin is reduced to resorufin, which produces very bright red fluorescence. Viable cells continuously convert resazurin to resorufin, thereby generating a quantitative measure of viability and cytotoxicity (Rampersad, 2012).

## 2. RESULTS

### 2.1. Fatty acid and volatile compounds from *Pistacia lentiscus* L. fruits oil

*Pistacia lentiscus* fruits oils from artisanal and commercial samples are mainly constituted by oleic acid (18:1), palmitic acid (16:0), linoleic acid (18:2), and palmitoleic acid (16:1) with a 3/2/1/0.1 *ratio*, being the four main fatty acids presenting more than 90% of fatty acids in these oils (Tab 1 and 2). These results are in accordance with data reported in the literature for Tunisian and Algerian samples (Charef et al, 2008; Mezni et al, 2012; Trabelsi et al, 2012). However, two commercial samples show very different fatty acid profiles with inversed ratio of linoleic and oleic acids, revealing probable adulteration of these two commercial products, as higher ratio of linoleic acid can be a consequence of linoleic rich oil addition such a sunflower oil (Christopoulou, 2004). The results indicate that *Pistacia lentiscus* oils present a medium UFA/SFA *ratio* between 1.54 and 2.57. Mezni et al. reported higher UFA/SFA ratio ranging between 2.33 and 2.84 (Mezni et al, 2012). In addition, other reports showed similar higher ratios as well (Charef et al, 2012; Trabelsi et al. 2012). These differences can be explained by differences in oil extraction protocols and systems. In fact, Charef et al. used Soxhlet extraction using hexane and Trabelsi et al. used petroleum ether by means of a Soxhlet extraction; whereas artisanal samples from this study were produced using traditional mechanical oil extraction methods.

In order to identify specific markers of *Pistacia lentiscus* oil using GC-MS, identification of other volatiles after derivatization are also examined. Monoterpenes are identified from retention times Rt=5 min to Rt=15 min (Tab. 3). Major monoterpenes are identified as α-pinene, myrcene, and β-limonene, largely reported in *Pistacia lentiscus* fruits and oil (Mecharara-Idjeli et al, 2008; Wylli et al, 2006). Myrcene is the most abundant monoterpene representing more than 50% of detected monoterpenes. Monoterpenes’ fraction may represent from 2.37 to 20.43% of all the identified volatiles compounds in the artisanal samples. Regarding artisanal samples (1 - 8), sample 7 *(El Kala)* presented the highest monoterpene fraction representing 20.43% of the detected compounds, *versus* only 2.36% for sample 2 (*Akbou 2015*) (Tab. 3). For commercial samples 9 (*El Wafia*) and 10 (*Zazia*), the monoterpenes fraction is nearly absent or not significant (0,26%), which confirms eventual adulteration of the samples, whereas other commercial samples (11-14) have a monoterpene fraction similar to those of artisanal samples which confirm their authenticity. It is interesting to note that from Rt=15 min to Rt=20 min, some sesquiterpenes could be identified by in small amount.

**Table 3.**
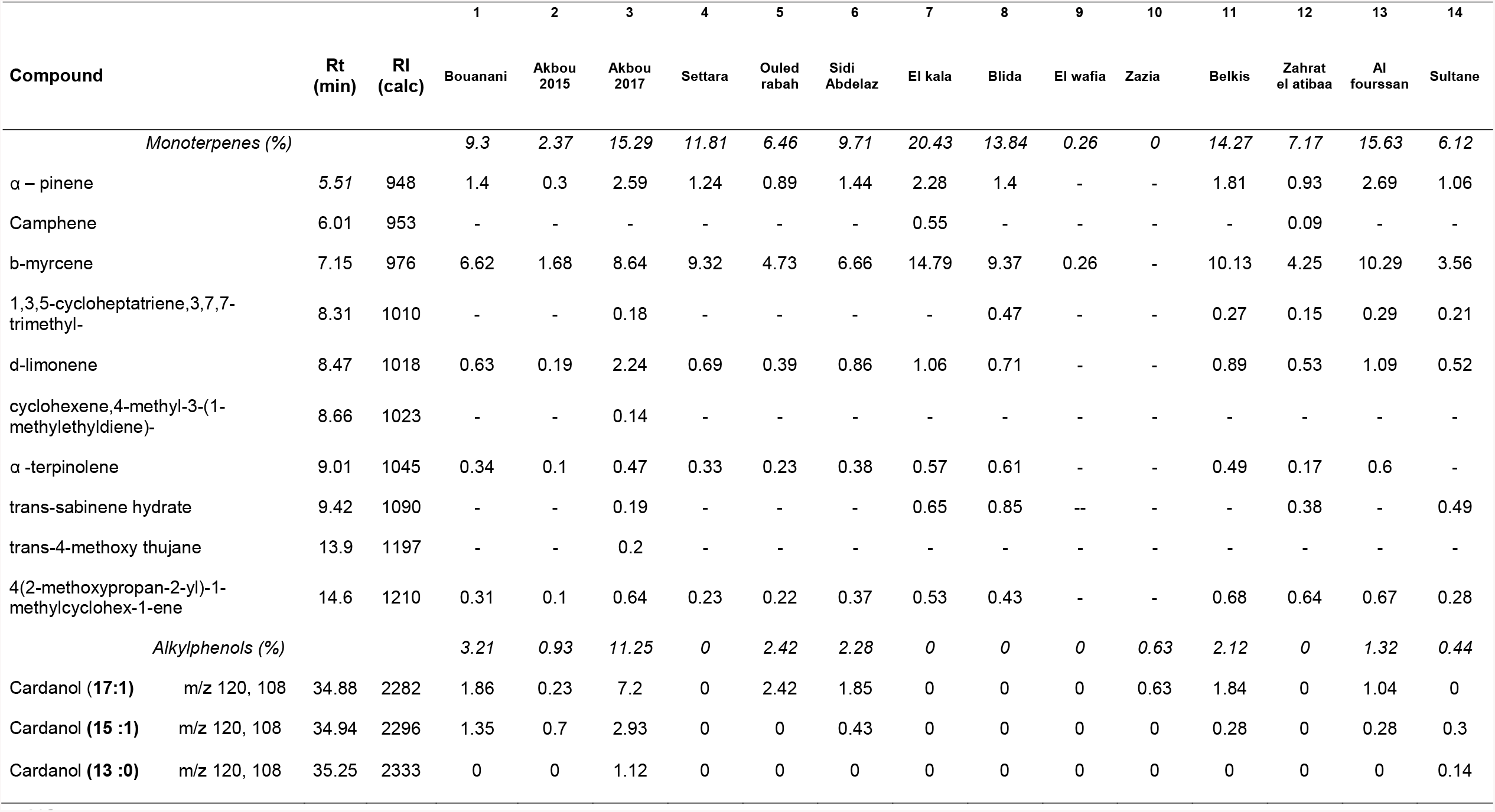
GC-MS identification of monoterpenes ans alkylphenols from Pistacia lentiscus oil.

After 34 min, traces of alkylphenols are identified using NIST library. The major MS fragments m/z value were 108 and 120. The literature report that GC/MS-based identification of cardanols from *Rhus* species describe the hydroxytropilium ion with a m/z value of 108 as a common fragment for all the identified alkylphenols (Frankie et al, 2001) (Tab. 3). Alkyl side chains for identifie compounds vary from 13 to 17 carbons and may present a probable unsaturation (Fig. 1). Occurrence of alkylphenols is considered a critical finding because of the irritating potential of these compounds in Anacardiaceae and Ginkgoaceae species.

**Figure 1.**
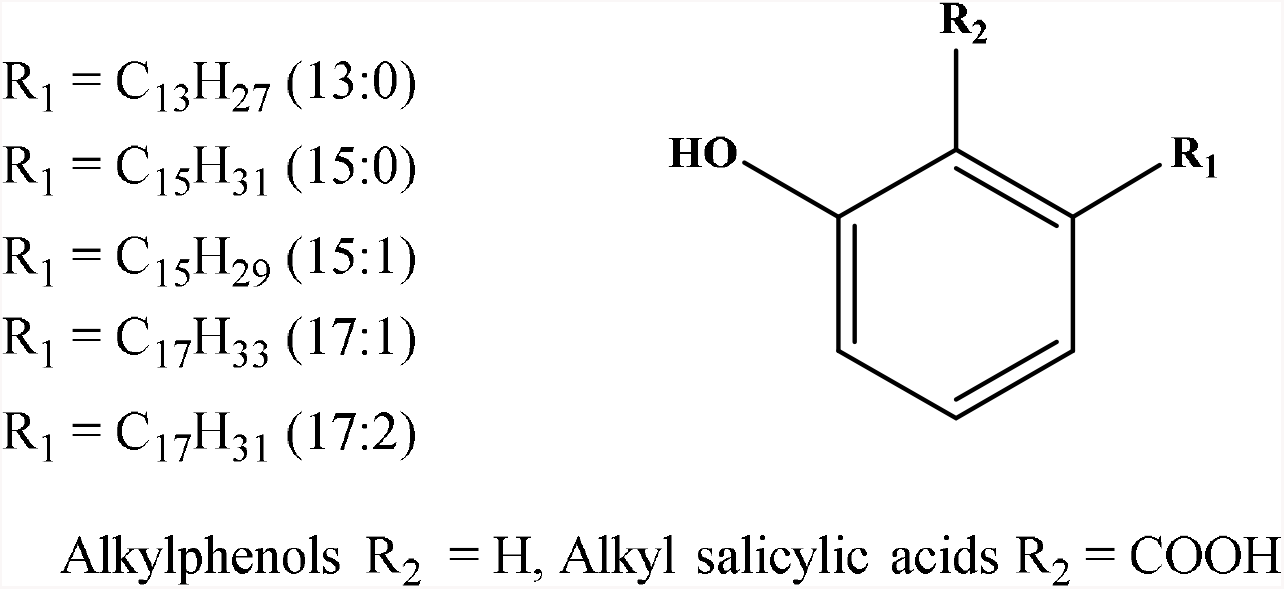
General structures of alkylphenols and alkylsalicylic acids

### 2.2. *Pistacia lentiscus* L. fruits oil unsaponifiable composition

Unsaponifiable extraction was considered for artisanal samples 2, 7, and 8 following AFNOR NF T 60-206. Composition of unsaponifiable fractions is analyzed using GC-MS after derivatization into silyl esters. Phytosterols and alkylsalicylic derivatives are identified using NIST library (Wang et al, 2014; Song et al, 2000). Five alkylsalicylic acids were identified between 11.5 and 14 min as C15:1, C15:0 and C17:1 derivatives. In this case, the diagnosis MS fragments was 180 (Tab. 4). In respect to phytosterols, identified between 26 and 36 minutes, β-sitosterol is the main identified phytosterol, stigmasterol and campesterol are also identified but in smaller proportions.

**Table 4.** GC-MS identification of phytosterols and alkylsalicylic acids from unsaponifiable.

| Ret.Time | m/z | Identification | 3b Akbou |  | 7b Elkala |  | 8b Blida |  |
| --- | --- | --- | --- | --- | --- | --- | --- | --- |
| *Alkylsalicylic acids silyl esters* |  |  | *Area* | *%* | *Area* | *%* | *Area* | *%* |
| 11.683 | 374, 207, 193, 180, 165 | C15:1 | 2021536 | 26.92 | 2910483 | 42.86 | 1416252 | 32.95 |
| 11.787 | 374, 207, 193, 180, 165 | C15:1 | 298418 | 3.97 | 294500 | 4.34 | 216470 | 5.04 |
| 11.851 | 376, 207, 193, 180, 165 | C15:0 | 699594 | 9.32 | 472092 | 6.95 | 310660 | 7.23 |
| 12.254 | 192, 143 | NI | 2071988 | 27.6 | 1325905 | 19.53 | 1620353 | 37.7 |
| 13.67 | 402, 180, 165 | C17:1 | 1480699 | 19.72 | 1092044 | 16.08 | 507068 | 11.8 |
| 13.765 | 402, 180, 165 | C17:1 | 936268 | 12.47 | 695181 | 10.24 | 227070 | 5.28 |
| *Phytosterols silyl esters* |  |  |  |  |  |  |  |  |
| 26.804 | 472, 382, 343, 207, 129 | Campesterol, TMS | 1008566 | 2.70 | 2728825 | 7.37 | 739221 | 3.0è |
| 27.174 | 483, 393, 207 | NI | 323628 | 0.87 | 257251 | 0.69 | 0 | 0 |
| 28.025 | 484, 394, 355, 281, 207 | Stigmasterol, TMS | 427520 | 1.14 | 1885985 | 5.10 | 265877 | 1.10 |
| 29.123 | 483, 393, 281, 207 | NI | 491899 | 1.32 | 381210 | 1.03 | 0 | 0 |
| 30.155 | 484, 394, 355, 281, 207 | NI | 193489 | 0.52 | 174070 | 0.47 | 0 | 0 |
| 30.476 | 486, 396, 381, 357, 255 | β-Sitosterol, TMS | 19841066 | 53.09 | 22865500 | 61.79 | 16281468 | 67.57 |
| 31.205 | 386, 281, 207 | NI | 1625896 | 4.35 | 1392926 | 3.76 | 834133 | 3.46 |
| 32.744 | 218, 189 | Lupeol TMS | 4269839 | 11.42 | 2826449 | 7.64 | 2239892 | 9.30 |
| 33.253 | 483, 393, 365, 339 | NI | 7122113 | 19.06 | 2761691 | 7.46 | 2148576 | 8.92 |
| 36.330 | 483, 422, 407, 379, 281 | NI | 2030622 | 5.43 | 1730922 | 4.68 | 1584655 | 6.58 |

### 2.3. Alkylsalicylic acids from Pistacia lentiscus L. fruits oil

As this is the first report demonstrating the occurence of alkylphenols or alkylsalicylic acids in *Pistacia lentiscus* L. oil, HPLC-DAD-MS analysis was envisaged with comparison of oil samples 1-14 to the 3-(heptadec-8-en-1-yl)-salicylic acid standard from PLFE1 fruit fraction. PLFE1 is a non-polar fraction rich in alkylsalicylic acids, is isolated from fruits of Algerian *Pistacia lentiscus* and characterized in a previous work (Tahrioui et al, 2020). PLFE1 contains 3-(heptadec-8-en-1-yl)-salicylic acid, also known as ginkgolic acid (C17:1), together with hydroginkgolic acid (C15:0). The structure of Δ8 ginkgolic acid (C17:1) have been confirmed after isolation from the PLFE1 extract using NMR analysis, whereas the double bound position was confirmed by ozonolysis (Tahrioui et al, 2020).

From HPLC-UV/DAD-MS analysis, similarity with UV spectra and m/z values in negative mode confirmed the identification of previously isolated alkylsalicylic acids in the non-polar fruit extract PLFE1. Figure 2 shows that alkylsalicylic acids derivatives are detected in oil samples between 44 and 60 minutes with the same distribution profile for 4 main compounds. Among the four detected derivatives, the main compounds were ginkgolic acid (C17:1) with a m/z value of 373 in negative mode, together with ginkgolic acid (C15:1) (m/z 345). The C17:2 derivative and the minor C13:0 derivative are also identified (m/z respectively 370.9 and 319) with similar typical UV spectra with two characteristic λmax at 247 and 314 nm (Fig.2).

**Figure 2.**
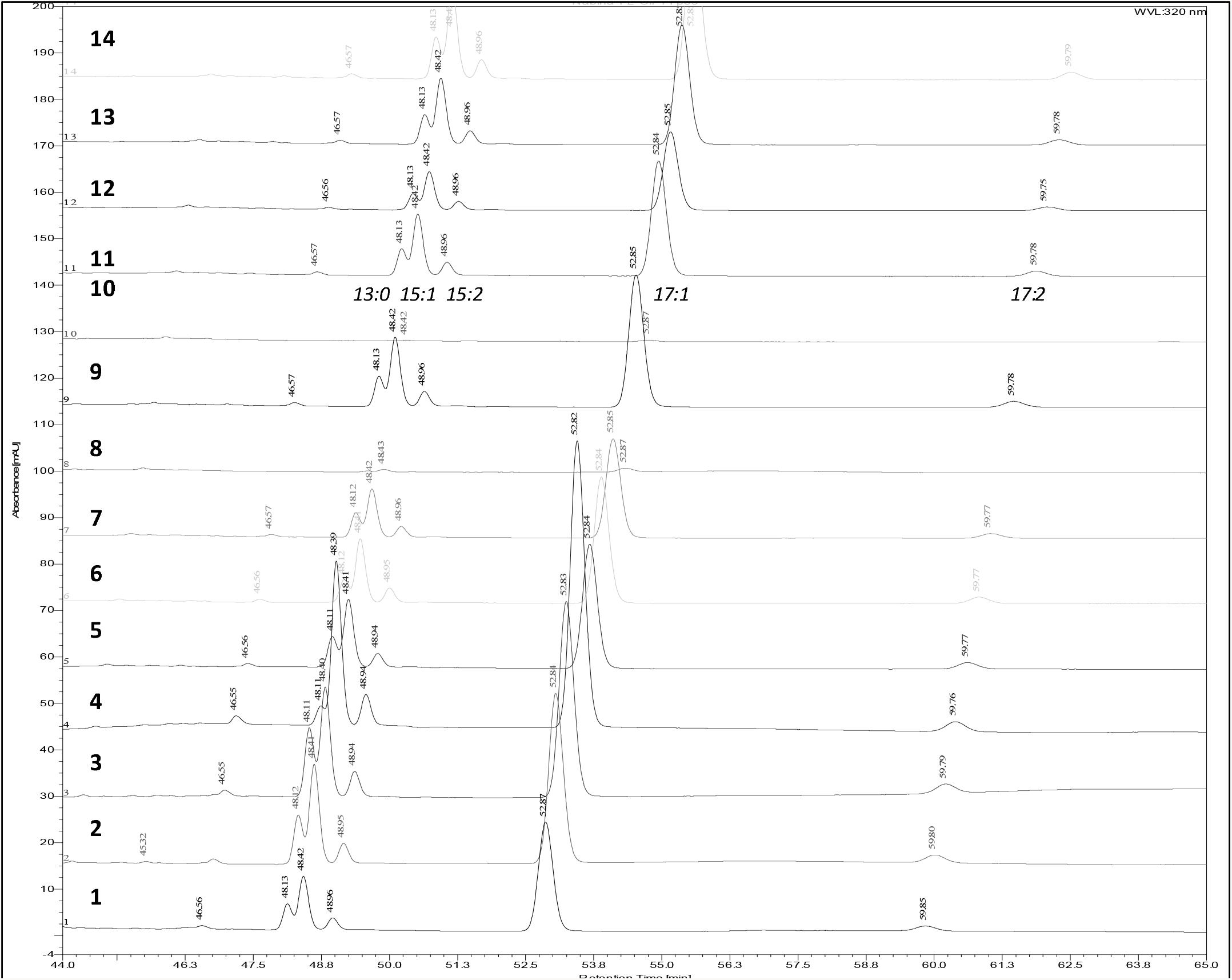
HPLC-DAD-MS profiles of *Pistacia lentiscus* oil fractions (44 to 65 minutes) showing the 5 main identified alkylsalicylic acids : C 13:0, C 15:1, C 15:2, C 17:1 and C 17:2.

### 2.4. Cytotoxic assessment on Normal Human Dermal Fibroblasts (NHDF)

Cell viability assay performed in NHDF cells revealed only a low cellular toxicity of the oil samples in all concentrations tested (0.1 to 100µg/mL), however, we detected a loss of cell viability for all unsaponifiable samples at concentrations higher than 50 µg/mL. At 100 µg/mL, cell viability for all unsaponifiable tested samples were below 50% (Fig. 3). These results may be linked to the presence of identified metabolites in the unsaponifiable. Regarding the different identified metabolites (phytosterols, alkylsalicylic acids, carotenoids), alkyl phenols and alkylsalicylic may be the highest contributors to this cytotoxicity as dermal toxicity of these class of compounds has already been described, such as contact dermatitis.

**Figure 3.**
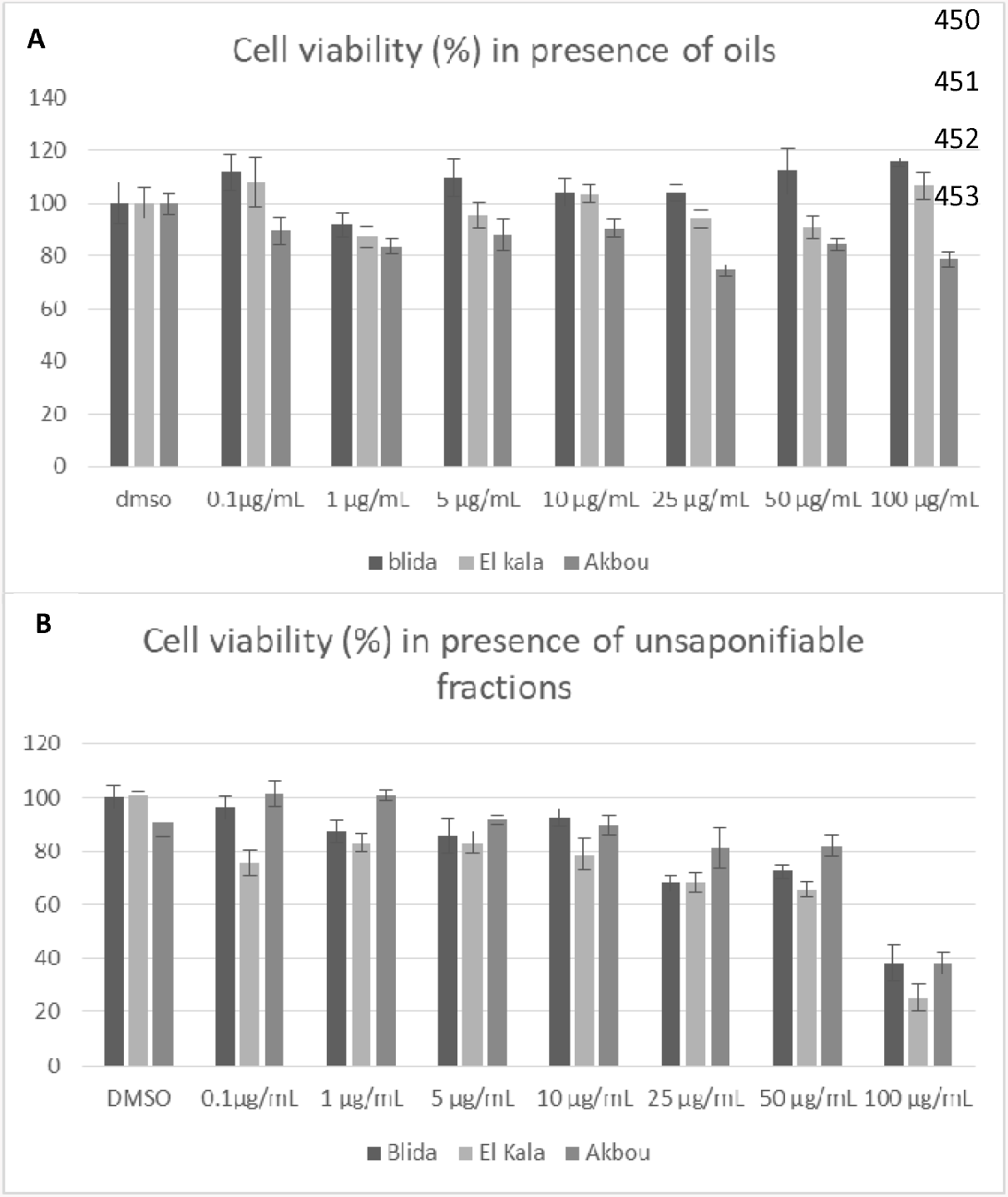
% of Cell viability (mean +: - SEM) of Normal Human Dermal Fibroblasts submitted to. (A) : *Pistacia lentiscus* L. artisanal oil samples 3 (Akbou 2017), 7 (El Kala) and 8 (Blida) from 0.1 to 100µg/mL, (B) Unsaponifiable fraction of *Pistacia lentiscus* L. artisanal oil 3b (Akbou 2017), 7b (El Kala) and 8b (Blida) from 0.1 to 100µg/mL.

## 3. DISCUSSION

Studied artisanal and commercial samples of *Pistacia lentiscus* oils were quite homogeneous in terms of fatty acids composition with oleic acid (18:1), palmitic acid (16:0), linoleic acid (18:2), and palmitoleic acid (16:1) profiling a 3/2/1/0.1 *ratio*, although two of the seven commercial samples are found to be adulterated (Tab. 1 and 2). Differences in trace secondary metabolites helped us identify alkylphenols derivatives as quality markers for *Pistacia lentiscus* fruit oils. HPLC-UV-DAD-MS confirmed that these alkylphenols are in fact alkylsalicylic acids after comparison with purified standards (Fig. 2). The 17:1 alkyl salicylic acid derivative is found to be the major derivative together with the 15:1 derivative. To the best of our knowledge, this is the first study to report the occurrence of alkylsalicylic acids (ASAs) in *Pistacia lentiscus* fruit oil. Previous identification of ASA in *Pistacia lentiscus* included previous authors findings of these ASAs in the non-polar fraction of the fruits particularly the 3-(heptadec-8-en-1-yl)-salicylic acid also known as ginkgolic acid (Δ8 C17:1) where the position of the double bound is determined using ozonolyisis (Tahrioui et al, 2020). This class of secondary metabolites is reported in other Anacadiaceae species, such as *Anacardium occidentale*, but also in Gingkoaceae species. Identification of alkylsalicylic acids by LC-MS and GC-MS is mainly described for *Ginkgo biloba* as quality requirements of phytopharmaceutical products requires limited amounts of ginkgolic acids (Abate-Pella, 2017). Alkylsalicylic acids could be identified in *Ginkgo biloba* leaves by GC-MS thanks to the characteristic fragmentation pattern with a typical fragment with m/z value of 180 described by Wang et al. for methyl esters derivatives (Wang et al, 2014). Alkylsalicylic acids’ profile in *Ginkgo biloba* leaves are quite different as the 15:1 ginkgolic acid, either Δ8 or Δ10, is the major derivative. Wang et al. determined the Δ8/Δ10 ratio using commercial standards on a HP-88 high polarity column, as the Δ8 compound retention time is slightly lower. In this study, GC-MS is performed with a RTx-5 non polar column. Indeed, two isomers of the 15:1 and 17:1 alkylsalicylic derivatives could be identified, without clear identification of each isomer. The availability of the purified standard fraction PLFE1 allowed us the identification of Δ8 17:1 derivative as the major alkyl salicylic acid compound present in *Pistacia lentiscus* fruit oil.

Alkylsalicylic acids, such as ginkgolic or anacardic acids, have been described as toxic compounds responsible for cutaneous irritation or allergy (Kajiyama et al, 2002; Njoko et al, 2000). However, this cutaneous toxicity observed after consumption of *Ginkgo biloba*, is still under scrutiny. Indeed, alkyl resorcinols, such as ginkgols and cardanols, or catechol derivatives, such as urushiols from Anacadiaceae species, are highly toxic as the resorcinol and catechol moiety might be transformed into quinones responsible of severe contact dermatitis (Duthil, 2005; Aguilar-Ortigoza, 2003; Knight et al, 1996). Alkylphenols and alkylresorcinols have also been identified in *Ginkgo biloba* leaves and may be responsible for the observed toxicity. Regarding alkylsalicylic acids derivatives, with a single phenol function, it is not clear yet whether transformation into quinones might occur after metabolization or not. Baron Ruppert and Luepke gave some evidence of cytotoxicity of a ginkgolic acids (GA) rich fraction (16% GA only) using the hen’s egg test. The low level of GA in the tested fraction is still not sufficient to explain the toxicity of GA (Lomonaco et al, 2013). However, more recently, hepatotoxicity of pure ginkgolic acids have been reported in mice and rat models (Baron-Ruppert et al, 2001). In the present work, evaluation of cytotoxicity on normal human dermal fibroblasts (NHDF) revealed that the unsaponifiable fraction can affect cell viability in fibroblasts, but this effect is not recorded for the oil (Fig. 3). While unsaponifiable fraction is rich in phyosterols and alkylsalicylic acids, toxicity assessment of phtoysterols in cosmetics revealed that they are usually not toxic (Jiang et al, 2017). Nevertheless, GA is considered a promising antitumor compound via the inhibition of the small ubiquitin-related modifiers SUMO-1 (Belsito et al, 2013) and as an antibacterial agent by inhibiting virulence factors such as biofilm formation or membrane stiffness (Tahrioui et al, 2020).

## 4. CONCLUSION

This work concerned the investigation of quality standards for *Pistacia lentiscus* fruits oil in commercial and artisanal samples lead to the identification of alkylsalicylic acids in fruits oil and unsaponifiable fraction of *Pistacia lentiscus* fruit oil. This is the first report of occurrence of alkylsalicylic acids in *Pistacia lentiscus* fruits oil. As alkylsalicylic acids are skin irritating agents, cytotoxicity evaluation on normal dermal human fibroblasts indicated that the oil is not toxic even at high concentrations, whereas unsaponifiable fraction containing higher amounts of alkylsalicylic derivatives are found to be toxic at concentrations above 50 µg/mL. As quality standards required for *Gingko biloba* phytopharmaceutical products is fixed by a limit of 5 ppm for alkylsalicylic acids is set; therefore, quantification of alkylsalicylic acids in *Pistacia lentiscus* oil both for nutraceutical or cosmeceutical use should be envisaged by manufacturers.

## Supporting information

Supplementary data Benalia 2020

## ACKNOWLEDGMENTS

Authors thank the funding program H2020 MSCA-RISE EXANDAS - EXploitation of Aromatic plaNts’ by proDucts for the development of novel cosmeceuticAls and food Supplements - H2020-MSCA-RISE-2015-grant agreement No 691247”. https://www.exandas-project.eu/.

## AUTHOR CONTRIBUTIONS

Conceptualization, methodology, S.B.; resources, S.B., N.B., S.C., formal analysis, S.B., N.B., A.B., S.O., T.R, B.L.F., investigation, S.B., N.B., A.B., S.O., E.C., T.R., data curation, N.B., S.B., A.B., S.O., E.C., T.R., writing—original draft preparation, S.B, N.B., T.R.; writing—review and editing, S.B., N.B., B.F., S.C., supervision, S.B., B.D., S.D., B.F.; project administration, S.B., B.D., S.D.; funding acquisition, S.B., B.D., S.M., S.D., S.C., B.F.

