## Supplementary data Benalia 2020 for "Identification of alkylsalicylic acids in Lentisk oil (*Pistacia lentiscus* L.) and cytotoxicity on Human Normal Dermal Fibroblasts": Benalia Supporting informations Biorvix.docx

**Supplementary material**

**Authors**

Nabiha Benalia^1,2^, Abdenour Boumechhour^3^, Sergio Ortiz^1^, Cristian Alejandro Echague^1^, Thorsten Rose^4^, Bernd L. Fiebich^4^, Smain Chemat^3^, Sylvie Michel^1^, Brigitte Deguin^1^, Saliha Dahamna^2^, Sabrina Boutefnouchet^1^*

**Affiliations**

1.	Université de Paris, UMR CNRS CiTCoM 8038, Faculté de Santé ; 4, av. de l’Observatoire, 75006 Paris, FRANCE.

2. Laboratory of Phytotherapy Applied to Chronic Diseases, Faculty of Natural and Life sciences, University Ferhat Abbas Setif 1, Setif 19000, ALGERIA..

3	Extraction and Separation Techniques Team, Research Centre in Physical and Chemical Analysis (C.R.A.P.C), BP 384 Zone Industrielle de Bousmail, CP 42004 Tipaza (ALGERIA) ;

4	VivaCEll Biotechnology GmbH, Ferdinand-Porsche-Str. 5, D-79211 Denzlingen, Germany

**Corresponding author(s)**

Sabrina Boutefnouchet^1 :^, +33253739811

**Abstract**

This data includes phytochemical analysis of *Pistacia lentiscus* fruits oil from Algeria obtained from samples of eight (08) artisanal and commercial (06) origin. These data include HPLC-DAD-MS based identification of alkylsalicylic acids in fruits oil and apolar fraction of fruits extract PLFE1, together with GC-MS identification of alkylsalicylic acids from unsaponifiable fraction. Spectroscopic data are also available relative to NMR characterization of the heptadecen-8-yl salicylic acid internal standard, isolated from PLFE1 fraction.

1. **Vegetal Oil samples**

Artisanal and commercial oil samples were collected in 2015 and 2017 and kept at +4°C for further analysis.

Material Transfert Agreement from Algeria to France available – EXANDAS Program : Grant Agreement number: 691247 — EXANDAS — H2020-MSCA-RISE-2015/H2020-MSCA-RISE-2015. <https://www.exandas-project.eu/>

Voucher specimen are available at : Université de Paris UMR 8038 – Faculté de Santé – UFR Pharmacie – 4, av. de l’Observatoire - Paris - France

Latitude and longitude and GPS coordinates for collected samples/data:

| **ID** | **Oil Sample** | **Area (a)** | **Nature of oil** | **GPS** |
| --- | --- | --- | --- | --- |
| 1 | Bouanani | Jijel | Semi-artisanal | 36° 43' 6.69" N, 6° 20' 9.244" E |
| 2 | Akbou 2015 | Bejaia | Semi-artisanal | 36° 27' 32.458" N, 4° 32' 0.522" E |
| 3 | Akbou 2017 | Bejaia | Semi-artisanal | 36° 27' 32.458" N, 4° 32' 0.522" E |
| 4 | Settara | Jijel | Artisanal | 36°42'05.1"N 6°20'01.5"E |
| 5 | Ouled rabeh | Jijel | Artisanal | 36°38'55.0"N 6°11'38.8"E |
| 6 | Sidi abdelaziz | Jijel | Artisanal | 36°50'43.8"N 6°03'08.8"E |
| 7 | El kala | El taref | Artisanal | 36°54'15.8"N 8°23'16.4"E |
| 8 | Blida | Blida | Artisanal | 36°28'29.5"N 2°51'12.9"E |
| 9 | El wafia | Blida | Commercial oil | - |
| 10 | Zazia | Oum bouaghi | Commercial oil | - |
| 11 | Belkis | Constantine | Commercial oil | - |
| 12 | Zahret atibaa | Skikda | Commercial oil | - |
| 13 | El fourssane | Setif | Commercial oil | - |
| 14 | Sultane | Setif | Commercial oil | - |

2. **Identification of Alkyl salicylic acids from Pistacia lentiscus fruits oil using HPLC-UV-DAD-MS Profiles**

Oil samples are analyzed on a HPLC-DAD-MS Thermo Scientific Dionex U3000 (Thermo-Dionex, Les Ulis, France) consisted of a quaternary pump (LPG-3400 SD), a thermostated autosampler (WPS-3000TSL), a thermostated column (TCC-3000SD), and a diode array detector (DAD-3000) on line with a quadrupole mass spectrometer (Surveyor MSQ plus System (Thermo-Dionex, Les Ulis, France). All oil samples were diluted in methanol (10 mg/mL). Solutions were filtered before injection on UptiDisc 0.45 M nylon filters (Interchim, Montluçon, France). 20 µL of each solution are injected and chromatograms are recorded at 210, 280, and 320 nm. Oven temperature is set at 30°C and the analysis is performed using a gradient elution: A (H_2_O, 0,5% formic acid) and B (ACN, 0.5% formic acid) as follows: 5% of B (0–10 min, isocratic), 5% to 100% of B (10-40 min, linear gradient), 100% of B (40-60 min, isocratic), 100% to 5% of B (60-70 min, linear gradient), 5% of B (70-75 isocratic) the flow rate is fixed at 0.5 mL/min.

2.1 **HPLC-UV-DAD-MS Profiles – Alkyl salicylic acids identification in oils samples**

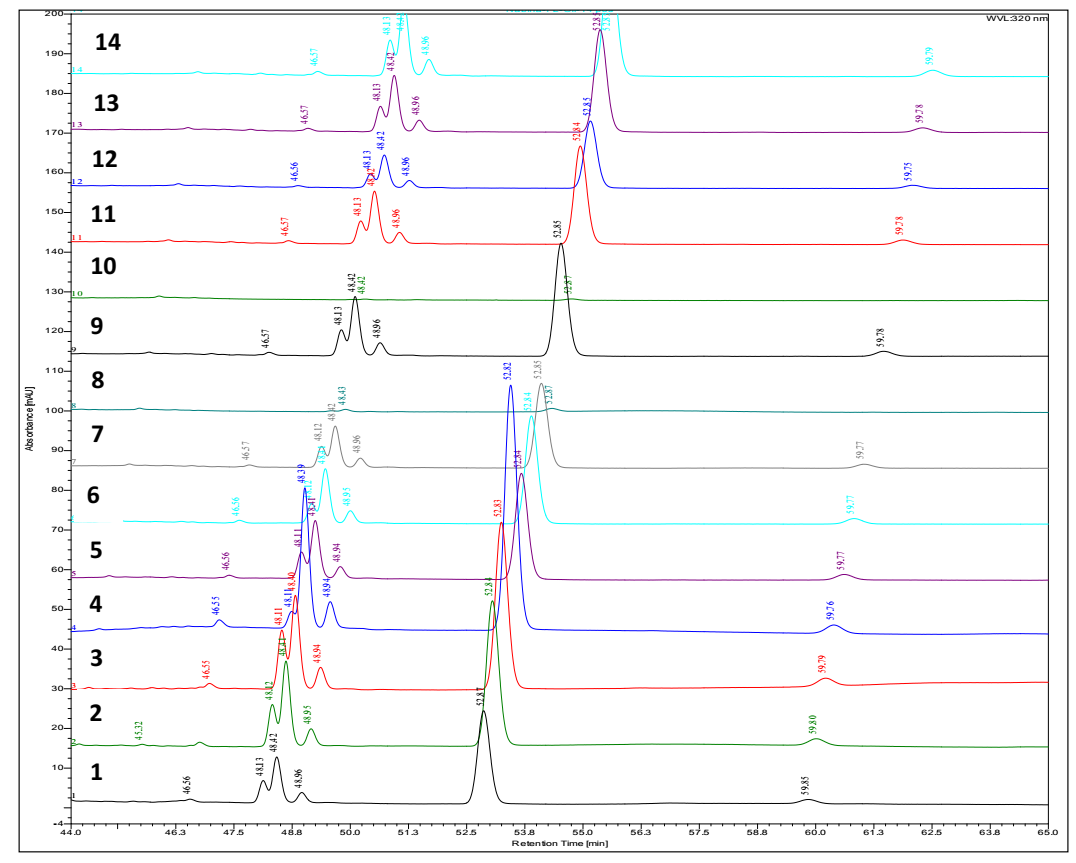

**17:0**

**17:1**

**15:1**

**13:0**

**15:2**

|  |  |  |  | PLFE1 | 1 | 2 | 3 | 4 | 5 | 6 | 7 | 8 | 9 | 10 | 11 | 12 | 13 | 14 |
| --- | --- | --- | --- | --- | --- | --- | --- | --- | --- | --- | --- | --- | --- | --- | --- | --- | --- | --- |
| RT min | SA | [M-H] m/z | λmax (nm) | Area% | | | | | | | | | | | | | | |
| 48.12 | 13 :0 | 319.00 | 246.5; 315.4 | 0.73 | 1.13 | 10.05 | 12.65 | 1.43 | 8.76 | 6.12 | 8.98 | - | 7.91 | - | 7.96 | 7.06 | 8.38 | 8.85 |
| **48.42** | **15 :1** | **345.01** | **246.7; 314.7** | **14.26** | **20.55** | **26.09** | **23.14** | **31.02** | **25.50** | **25.19** | **22.86** | **-** | **24.76** | **-** | **25.10** | **23.74** | **25.64** | **25.66** |
| 48.95 | 17 :2 | 370.94 | 246.8;314.8 | 6.94 | 5.25 | 6.30 | 6.87 | 6.09 | 5.97 | 6.52 | 6.13 | - | 6.20 | - | 6.08 | 6.06 | 5.90 | 6.24 |
| **52.83** | **17 :1** | **373.03** | **247.0;314.8** | **72.84** | **67.01** | **55.18** | **54.84** | **57.99** | **56.87** | **59.28** | **59.31** | **-** | **57.33** | **-** | **57.17** | **59.25** | **56.37** | **55.39** |

HPLC-UV-DAD-MS identification of alkylsalicylic acids identified in artisanal and commercial *Pistacia lentiscus* oil and PLFE1 apolar fruits extract.( % area of detected peaks between 45 and 60 minutes). SA : identified salicylic acid

2.2 **HPLC-UV-DAD-MS Profiles – Alkyl salicylic acids in oils samples: UV spectra and MS spectra**

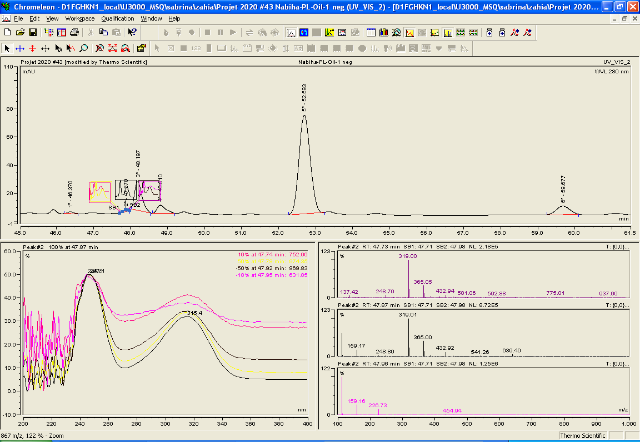

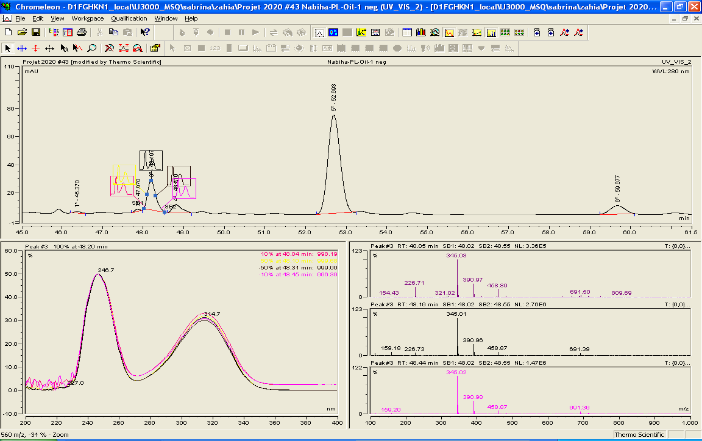

**13:0**

**15:1**

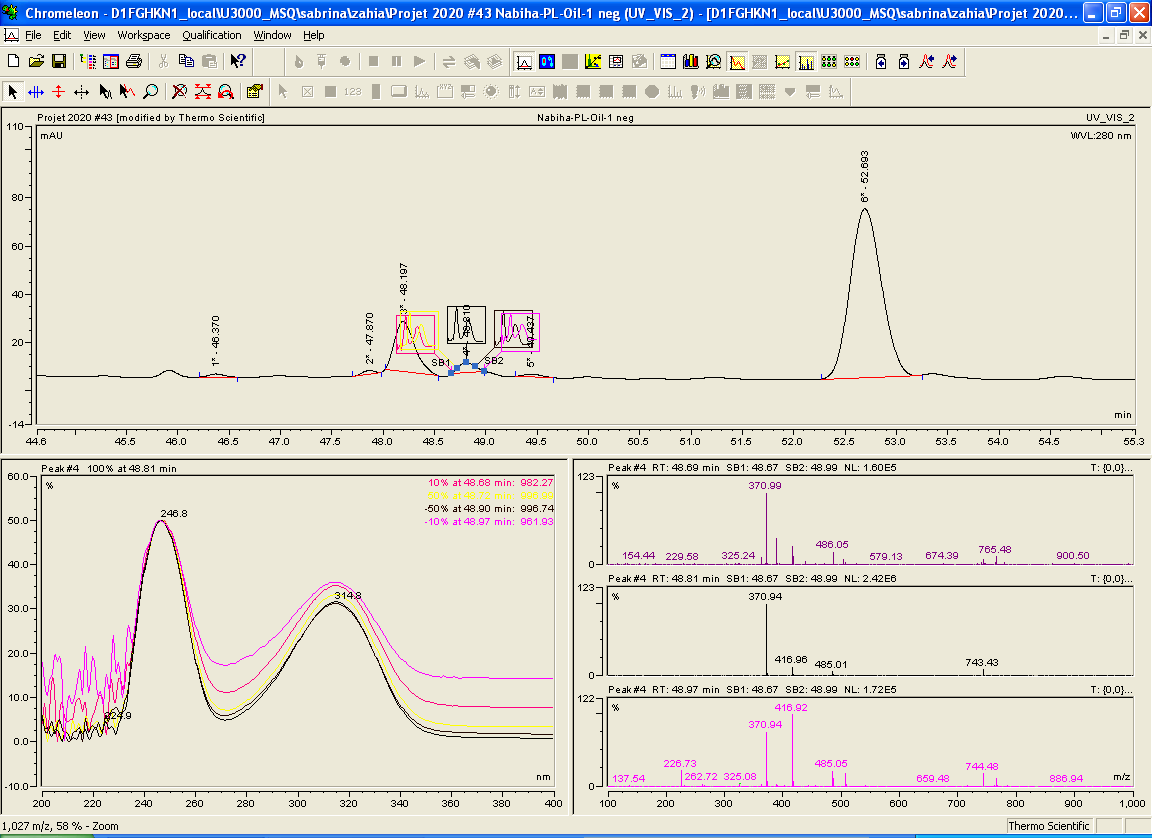

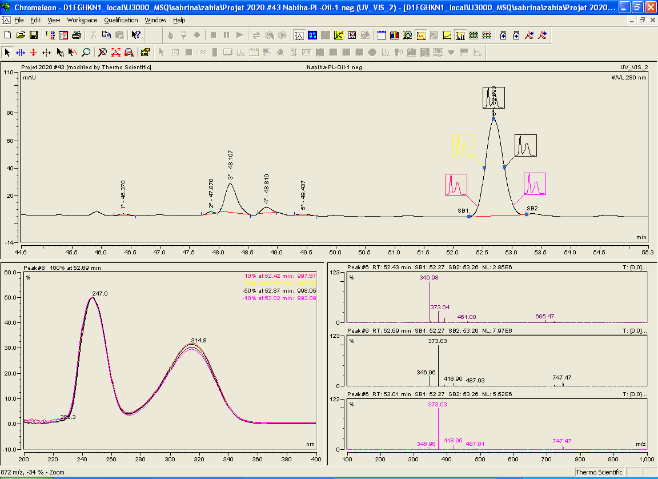

**17 :1**

**17:2**

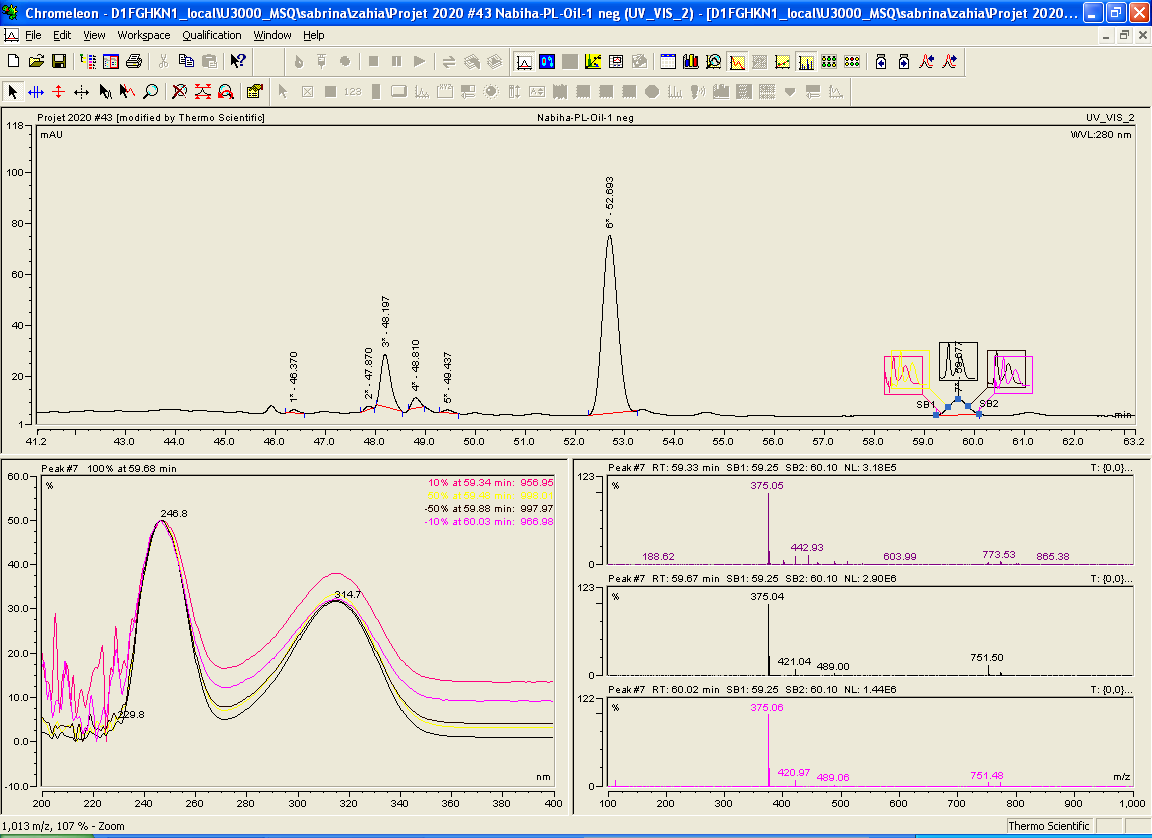

**17 :0**

**2.3 HPLC-UV-DAD-MS profile of PLFE1 apolar fraction**

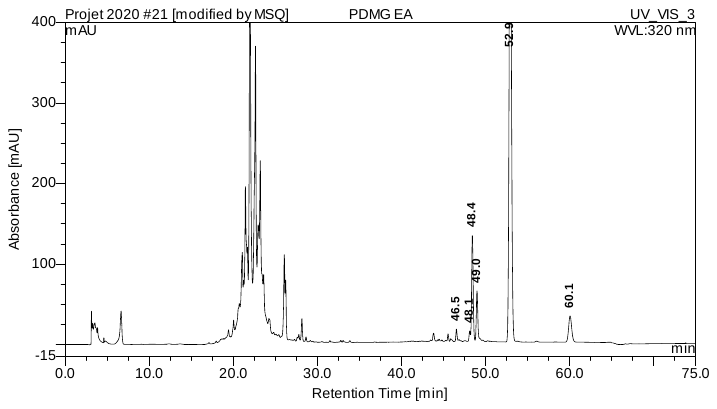

| **No.** | **Ret.Time** | **Peak Name** | **Height** | **Spectrum Name** | | **Lib.Name** | **Match** |
| --- | --- | --- | --- | --- | --- | --- | --- |
|  | **min** |  | **mAU** | **in Library** |  |  | **with Lib** |
| **1** | **46.54** | **n.a.** | **13.883** | **n.a.** |  | **n.a.** | **n.a.** |
| **2** | **48.13** | **n.a.** | **6.324** | **n.a.** |  | **n.a.** | **n.a.** |
| **3** | **48.44** | **n.a.** | **124.184** | **n.a.** |  | **n.a.** | **n.a.** |
| **4** | **48.99** | **n.a.** | **60.428** | **n.a.** |  | **n.a.** | **n.a.** |
| **5** | **52.94** | **n.a.** | **634.361** | **n.a.** |  | **n.a.** | **n.a.** |
| **6** | **60.06** | **n.a.** | **31.750** | **n.a.** |  | **n.a.** | **n.a.** |
| **Total:** |  |  | **870.931** |  |  |  | **n.a.** |

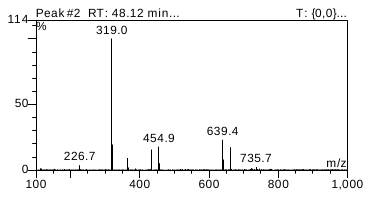

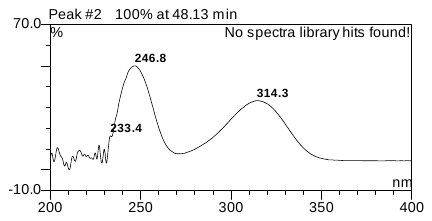

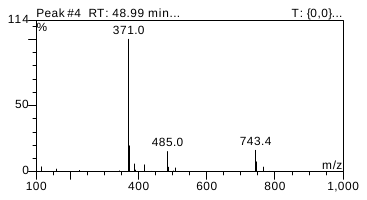

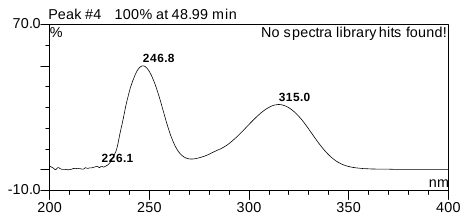

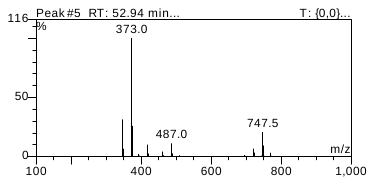

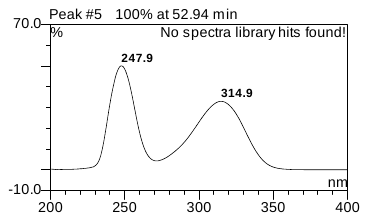

3. **Isolation from Pistacia lentiscus fruits and NMR structural characterization of 3-(heptadec-8-en-1-yl)-salicylic acid (17:1)**

***MPLC procedure*** *:* Stationary Phase: 9,6 g de Silica 60 M (0,04 – 0,063 mm), mobile phase : cyclohexane / ethyl acetate (gradient) , Flow : 5 - 10 mL/min, Pressure : 0,2 – 0,3 bar , Crude : 200 mg of PLFE-1 from *P.* *lentiscus. 35.5 mg of pure 3-(heptadec-8-en-1-yl)-salicylic acid (17:1).* To identify the double bond position, an ozonolysis reaction is carried out as described in earlier studies (Tahrioui, et al. ; Membrane-Interactive Compounds From *Pistacia lentiscus* L. Thwart *Pseudomonas aeruginosa* Virulence. *Frontiers in microbiology*, **2020**, *11*, 1068. https://doi.org/10.3389/fmicb.2020.01068 ). 1D and 2D NMR experiments are conducted in a Bruker 400 MHz spectrometer apparatus (Wissembourg, France). GCMS-QP2010 Ultra (Shimadzu Co. Kyoto. Japan) equipped with a fused silica capillary column (Rtx-5MS; 30 m × 0.25 mm inner diameter, film thickness 0.25 μm, Thames Restek. UK), and an Agilent 5973N MS detector.

**NMR spectra** obtained from solution of purified compound in deuterated chloroform (CDCl_3_), 1H NMR, 13C RMN, 1H – 1H – COSY , HSQC, HMBC were recorded**.**. Ozonolysis was performed on a dichloromethane solution of purified compound until red coloration appears. The ozonolysis product was analysed using GC-MS. The vector gas was Helium with a flow of 1.0 mL/min, split 1:20. The temperature program was as follows : initial oven temperature 70ºC during 5 minutes, then 120ºC (5ºC/min), 2 minutes step at 120°C then heat up until 180ºC (30ºC/min). After a 12 minutes step et 180°C, the temperature was heated up to 270ºC (30ºC/min) and hold during 20 minutes. Injection volume : 1 μL. Temperature of the injection chamber was 200ºC and temperature of the interface was kept at 290ºC.

### 3.1. Structural analysis of heptadecen-8-yl salicylic acid Δ8 - C17:1, isolated from PLFE1 *P. lentiscus* fruits apolar fraction

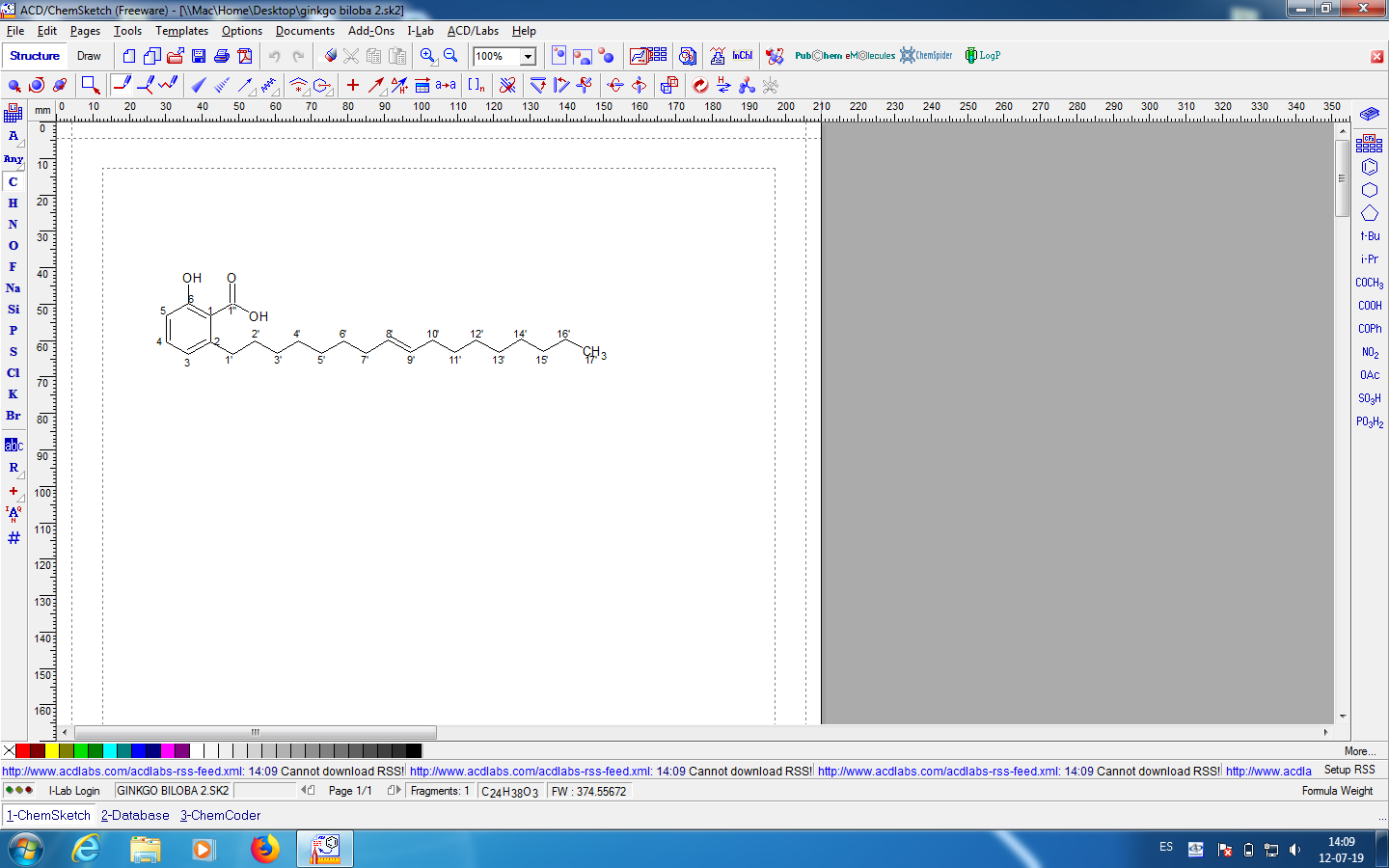

**^1^H NMR of Δ8 - C17:1 (solvent CDCl_3_).**

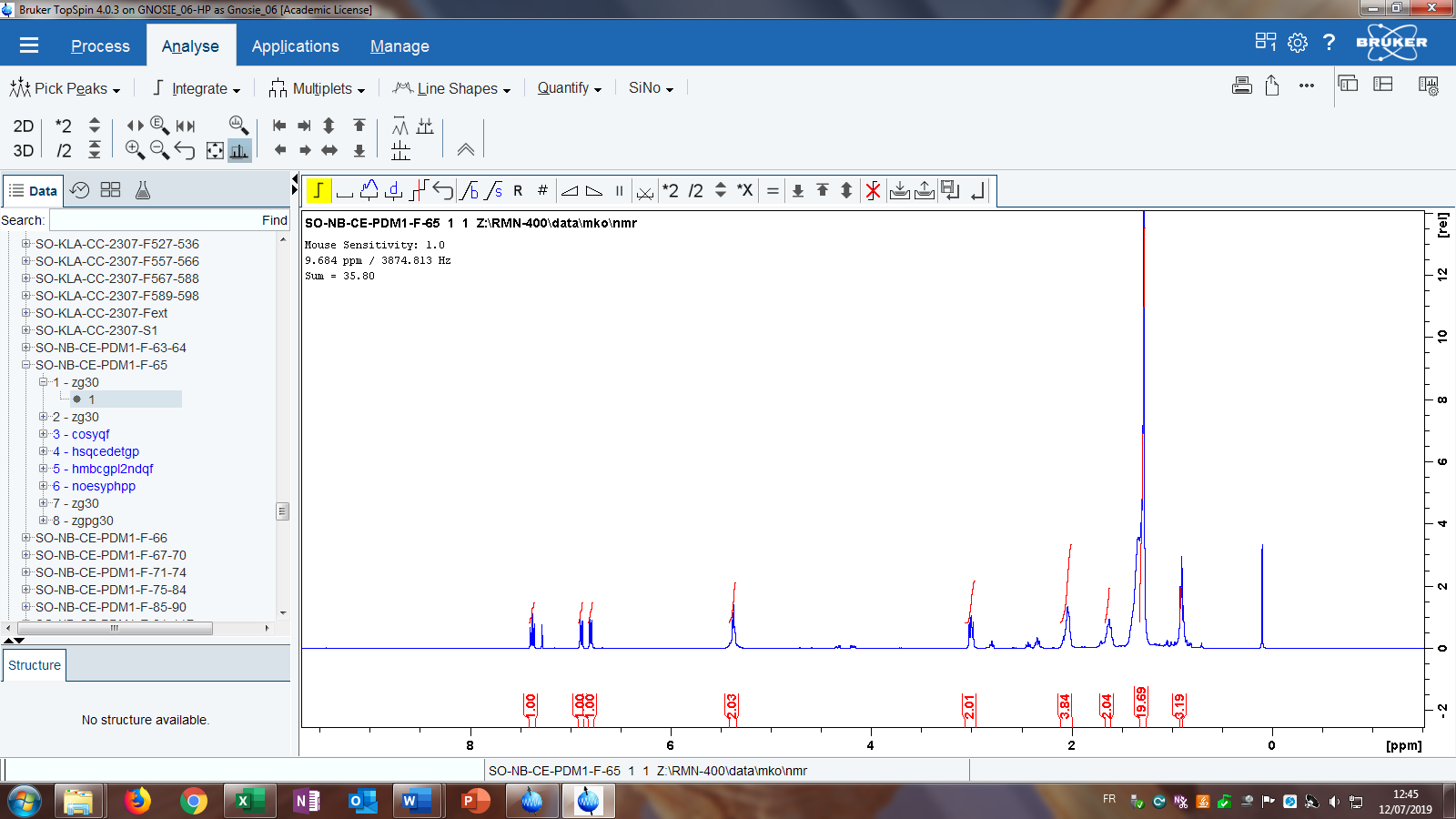

| **∂ (ppm)** | **Intégration** | **Multiplicité du signal** | ***J* (Hz)** | **Attribution** |
| --- | --- | --- | --- | --- |
| **7.36** | **1 H** | **Triplet** | **8.2** | **H4** |
| **6.88** | **1 H** | **Doublet** | **8.3** | **H5** |
| **6.78** | **1 H** | **Doublet** | **8.3** | **H3** |
| **5.36** | **2 H** | **Triplet** | **4.6** | **H8’, H9’** |
| **2.98** | **2 H** | **Triplet** | **7.7** | **H1’** |
| **2.04** | **4 H** | **Multiplet** |  | **H7’, H10’** |
| **1.63** | **2 H** | **Multiplet** |  | **H2’** |
| **1.28** | **20 H** | **Multiplet** |  | **H3’ – H6’, H11’ – H16’** |
| **0.88** | **3 H** | **Triplet** | **6.1** | **H17’** |

**^13^C RMN of Δ8 - C17:1,**

| **∂ (ppm)** | **Attribution** |
| --- | --- |
| **163.6** | **C 6** |
| **115.9** | **C 3** |
| **135.4** | **C 4** |
| **122.7** | **C 5** |
| **147.7** | **C 2** |
| **110.4** | **C 1** |
| **175.5** | **C 1’’** |
| **130.0 (*)** | **C 8’ (*)** |
| **129.8 (*)** | **C 9’ (*)** |
| **36.5** | **C 1’** |
| **32,0** | **C 2’** |
| **29.0** | **C 3’ – C 6’, C 11’ – C 16’** |
| **27.2 (**)** | **C 7’ (**)** |
| **26.9 (**)** | **C 10’ (**)** |
| **14.1** | **C 17’** |

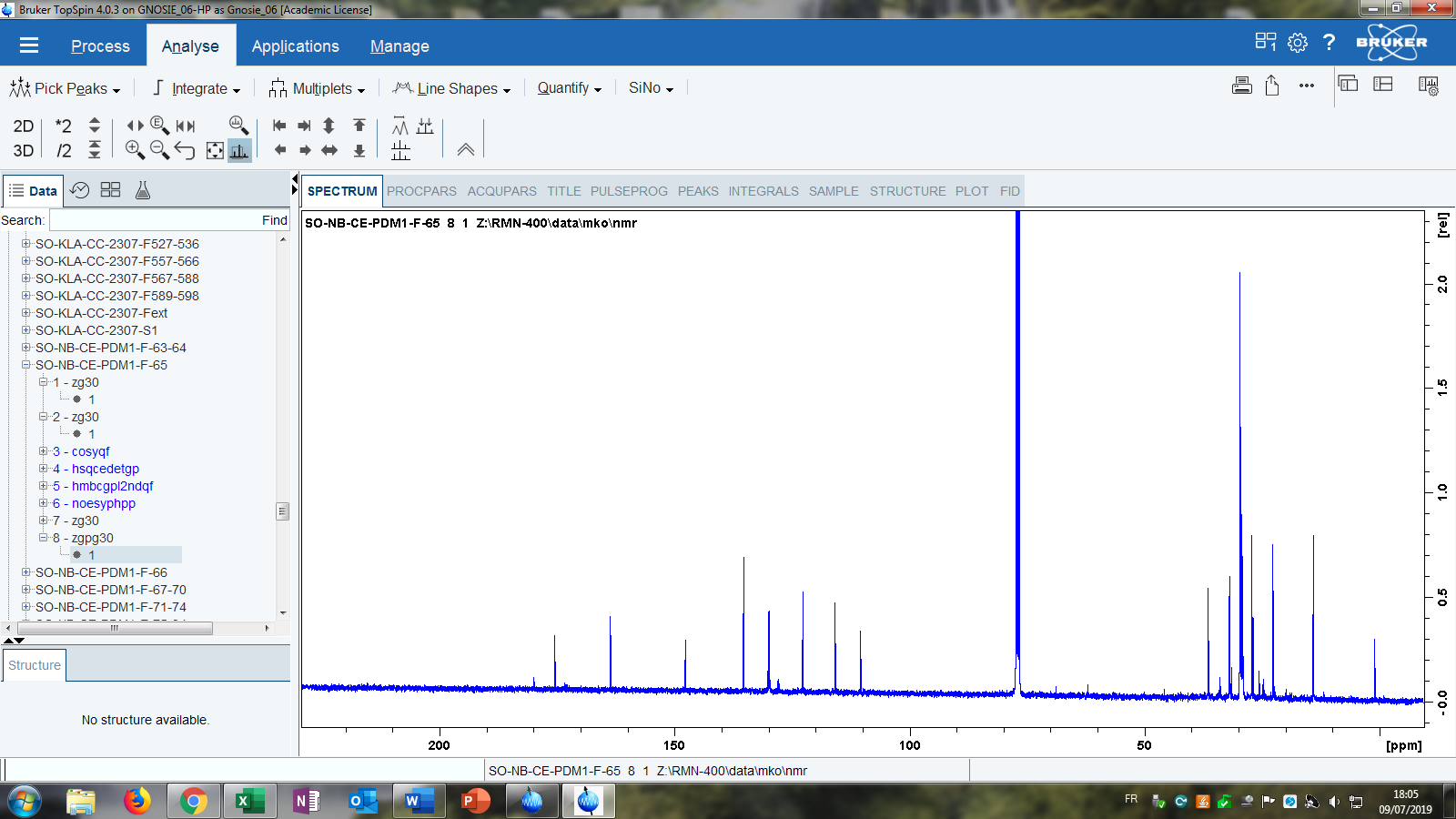
 **^13^C RMN of Δ8 - C17:1,**

**^1^H – ^1^H – COSY of Δ8 - C17:1,**

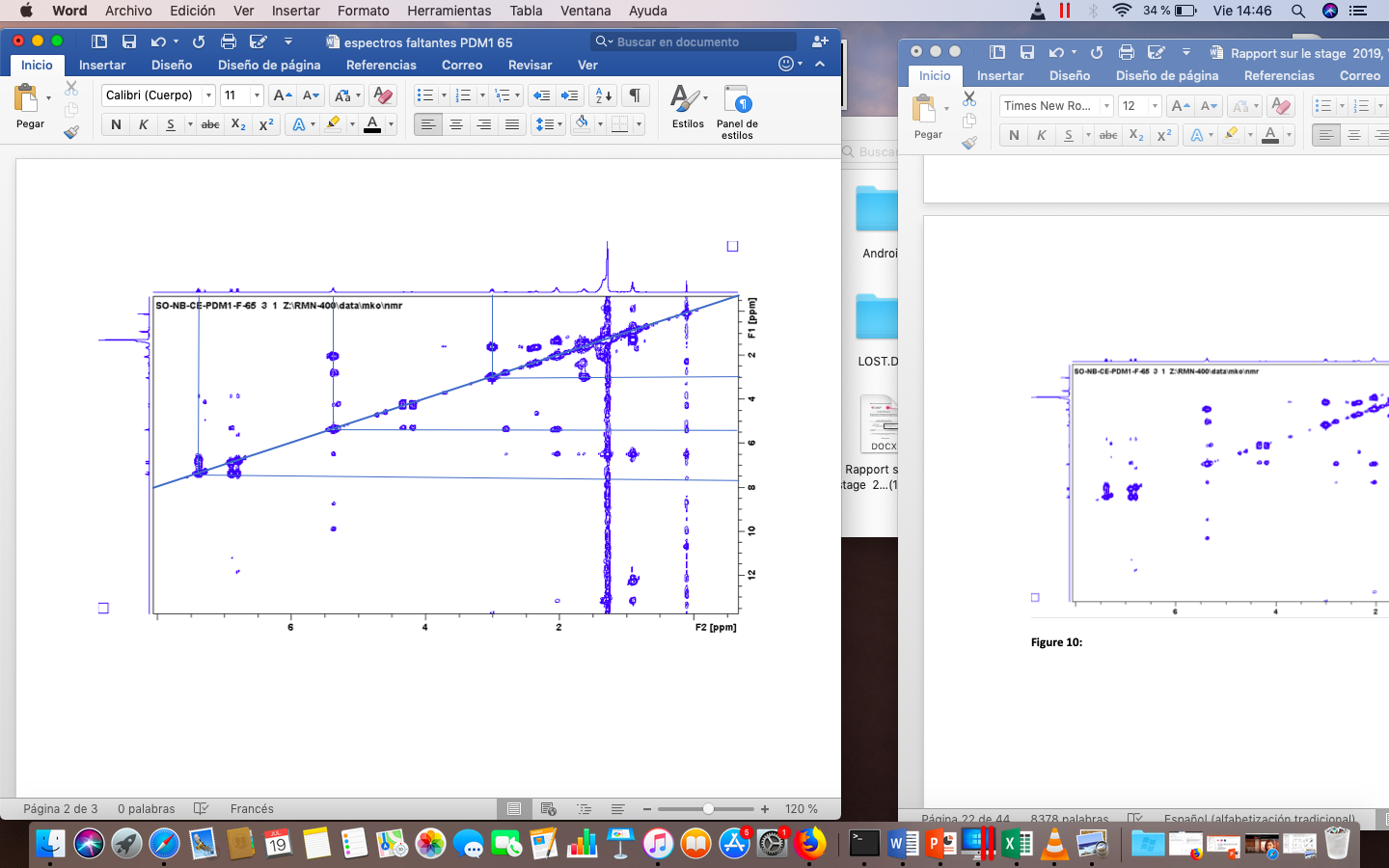

**HSQC of Δ8 - C17:1**

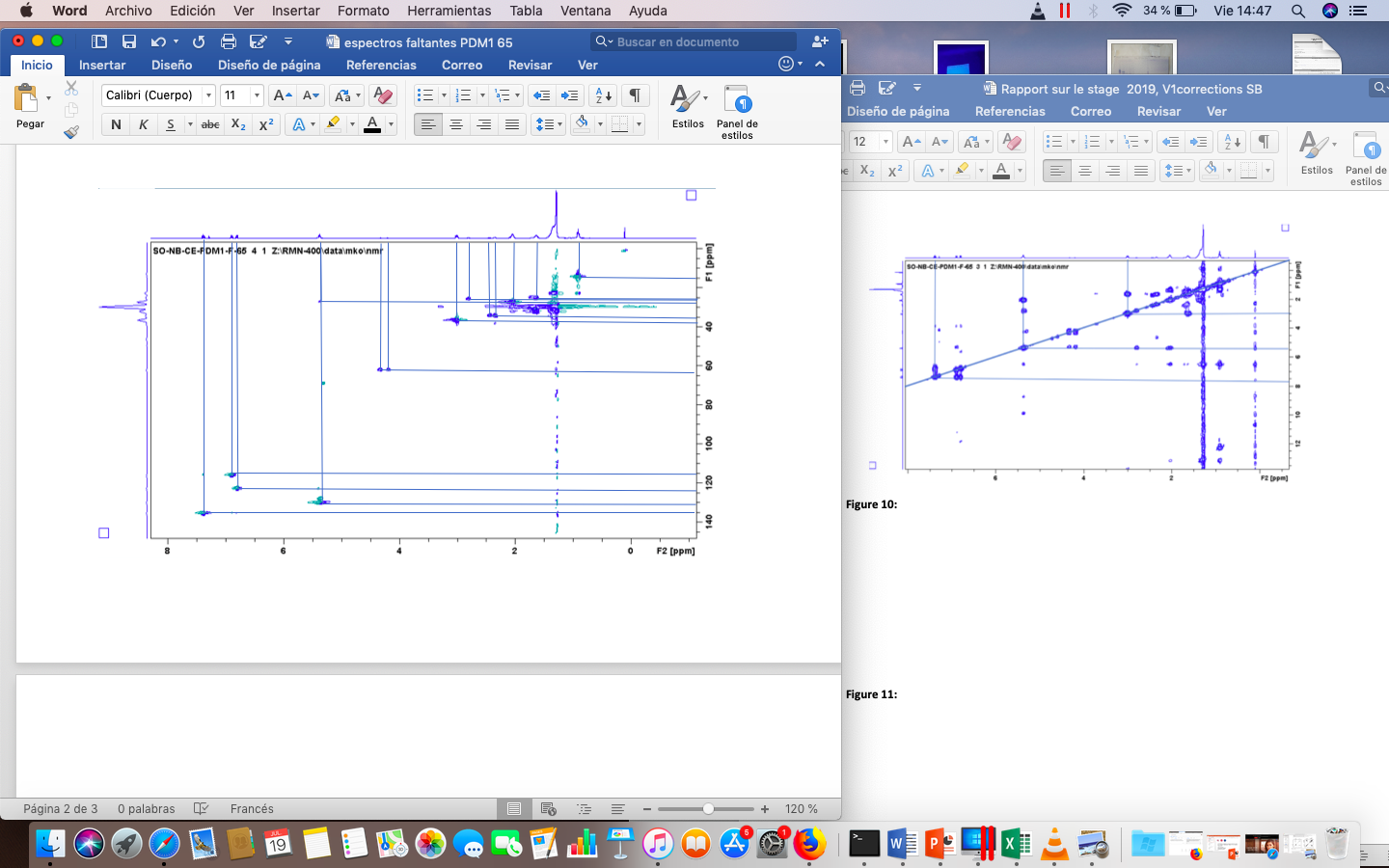

| **∂ (ppm)** | **Attribution** |
| --- | --- |
| **7.36 / 135.4** | **C4 – H4** |
| **6.88 / 115.9** | **C5 – H5** |
| **6.78 / 122.7** | **C3 – H3** |
| **5.36 / 130.0 , 5.36 / 129.8** | **C8’ - H8’ (*), C9’- H9’ (*)** |
| **2.98 / 36.5** | **C 1’ – H1’** |
| **2.04 / 27.2, 2.04 / 26.9** | **C7’ – H7’ (**), C10’- H10’ (**)** |
| **1.63 / 32.0** | **C2’ – H2’** |
| **1.28 / 29.0** | **C 3’-H3’ – C 6’-H6’, C11’-H11’ – C 16’-H16’** |
| **0.88 / 14.1** | **C17’ – H17’** |

**HMBC of Δ8 - C17:1,**

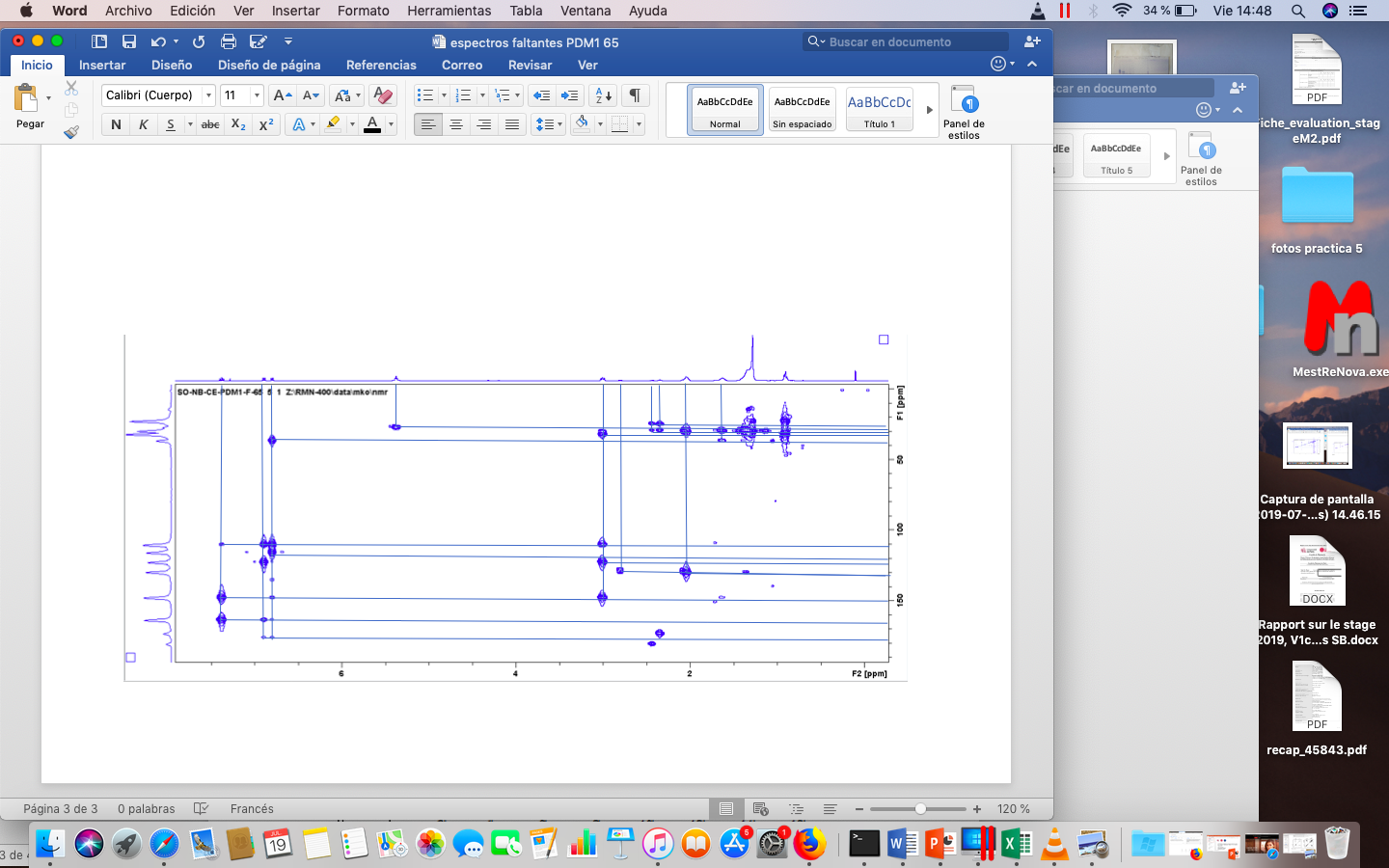

#### GC-MS analysis of products obtained from ozonolysis

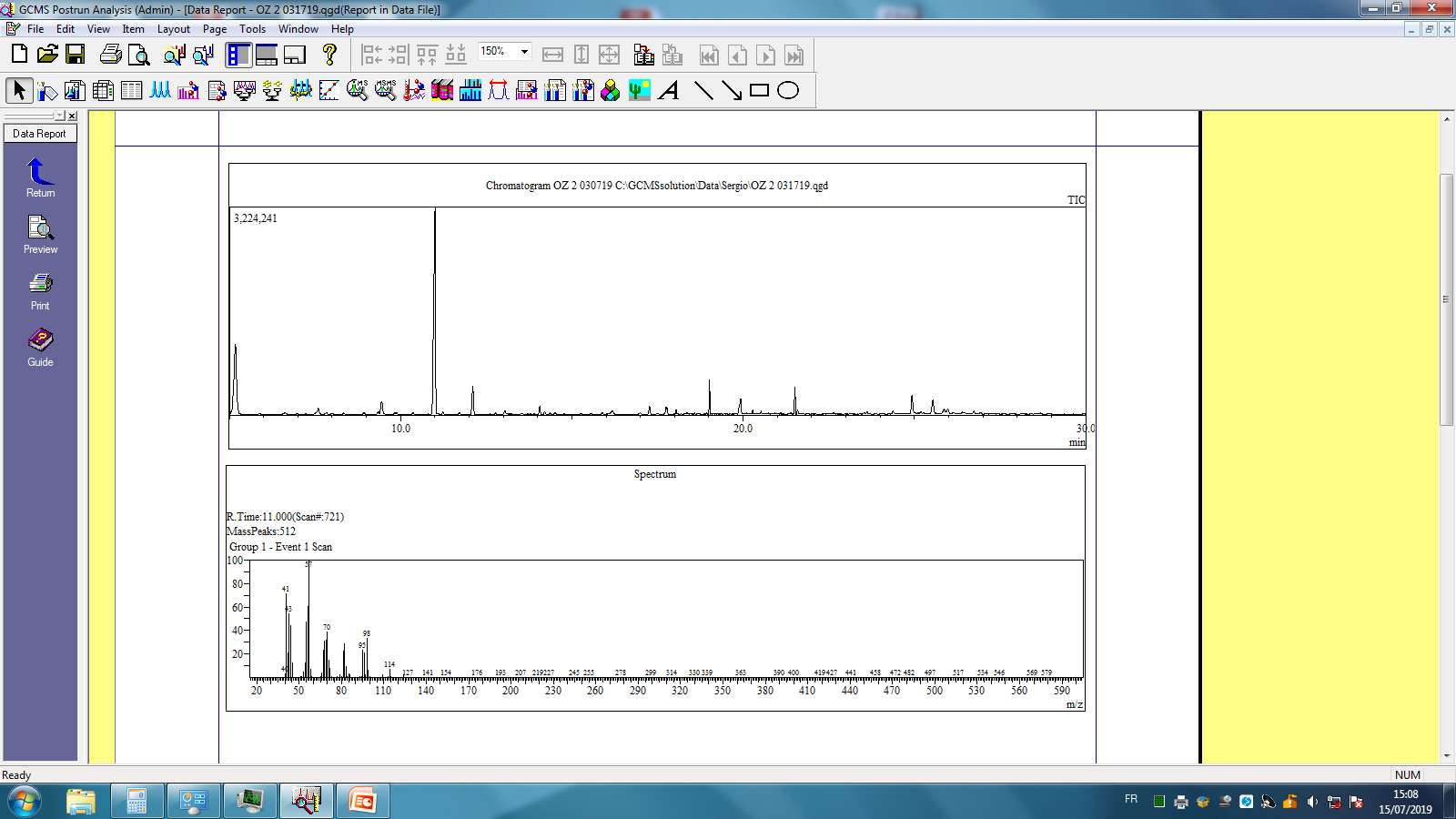

**MS chromatogram and MS spectra of main compound (RT 11 min).**

4. **Identification of alkylsalicylic acids in Pistacia lentiscus fruits oil unsaponifiable fractions using GC-MS (silylated derivatives)**

The recovery of unsaponifiable fraction for artisanal samples 2, 7, and 8 is conducted following the procedure described in AFNOR NF T 60-206. Briefly, 50 mL of 2N ethanolic solution of KOH are added to 5 g of oil. The mixture is then refluxed for one hour. After evaporation, 50 mL of water is added, the suspension is extracted three times with 100mL diethyl ether (3x100 mL), washed with aqueous KOH (0.5 N) followed by water, then dried on anhydrous Na_2_SO_4_ and evaporated under vacuum. Then, derivatization into silyl esters is performed. Briefly, 5 mg of unsaponifiable fraction are placed with 0.5 mL of pyridine in a 2 mL vial. Then, 0.1 mL of hexamethyldisilazane (HMDS) and 0.04 mL of trimethylchlorosilane (TMCS) are added and the reaction mixture is mixed using a vortex then centrifuged. From the supernatant of the silylated mixture, 1 µL is directly submitted to GC-MS analysis.

GCMS-QP2010 Ultra (Shimadzu Co. Kyoto. Japan) equipped with a fused silica capillary column (Rtx-5MS; 30 m × 0.25 mm inner diameter, film thickness 0.25 μm, Thames Restek. UK), and an Agilent 5973N MS detector. . In this case, carrier gas is H_2_ with a flowrate of 1 mL/min and a split 1:20, oven is programmed increasing from 180°C to 270°C at 8°C/min with a hold at initial and final temperatures of 1 and 65 min respectively [11]. MS Interface temperature is set at 240 °C while MS source temperature is set at 220 °C with an ionization energy of 70 eV. The injection volume was 1 µL.

**Alkylsalicylic acids in unsaponifiable fractions GC-MS (silylated derivatives)**

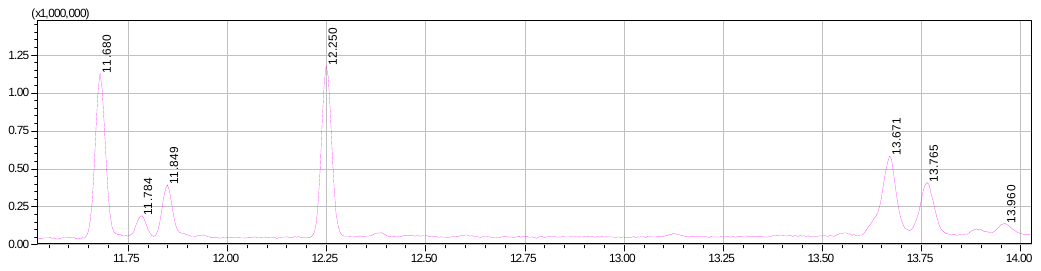

**17:1**

**17:1**

**15:0**

**15:1**

**15:1**

**3b-*Akbou***

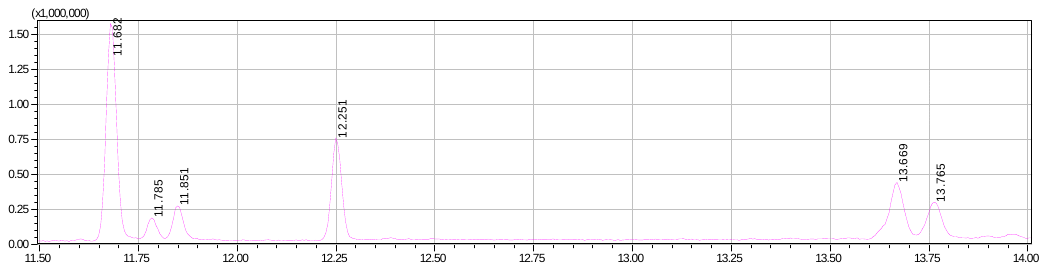

**15:1**

**15:1**

**15:0**

**17:1**

**17:1**

**7 *El Kala***

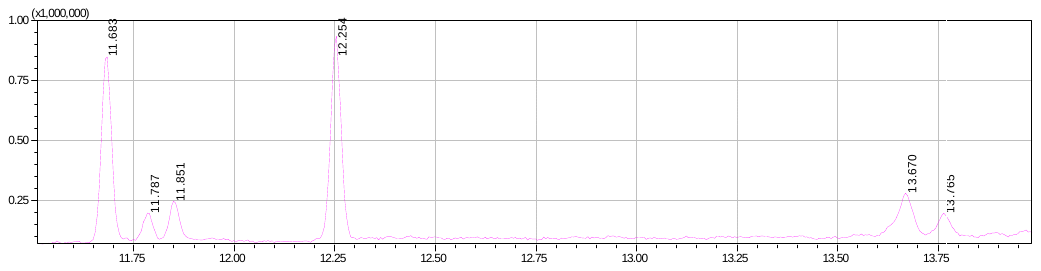

**17:1**

**17:1**

**15:0**

**15:1**

**15:1**

**8 *Blida***

| **Ret.Time** | **m/z** | **Identification** | **3b Akbou** |  | **7 Elkala** |  | **8 Blida** |  |
| --- | --- | --- | --- | --- | --- | --- | --- | --- |
| **Alkylsalicylic acids** | |  | **Area** | **%** | **Area** | **%** | **Area** | **%** |
| **11.683** | **180, 165** | **C15:1** | **2021536** | **26.92** | **2910483** | **42.86** | **1416252** | **32.95** |
| **11.787** | **180, 165** | **C15:1** | **298418** | **3.97** | **294500** | **4.34** | **216470** | **5.04** |
| **11.851** | **376, 180, 165** | **C15:0** | **699594** | **9.32** | **472092** | **6.95** | **310660** | **7.23** |
| **12.254** | **192, 143** | **NI** | **2071988** | **27.6** | **1325905** | **19.53** | **1620353** | **37.7** |
| **13.67** | **180, 165** | **C17:1** | **1480699** | **19.72** | **1092044** | **16.08** | **507068** | **11.8** |
| **13.765** | **180, 165** | **C17:1** | **936268** | **12.47** | **695181** | **10.24** | **227070** | **5.28** |
|  |  |  | **7508503** | **100** | **6790205** | **100** | **4297873** | **100** |
